# AMPlify: attentive deep learning model for discovery of novel antimicrobial peptides effective against WHO priority pathogens

**DOI:** 10.1101/2020.06.16.155705

**Authors:** Chenkai Li, Darcy Sutherland, S. Austin Hammond, Chen Yang, Figali Taho, Lauren Bergman, Simon Houston, René L. Warren, Titus Wong, Linda M.N. Hoang, Caroline E. Cameron, Caren C. Helbing, Inanc Birol

## Abstract

Antibiotic resistance is a growing global health concern prompting researchers to seek alternatives to conventional antibiotics. Antimicrobial peptides (AMPs) are emerging therapeutic agents with promising utility in this domain and using *in silico* methods to discover novel AMPs is a strategy that is gaining interest. Such methods can filter through large volumes of candidate sequences and reduce lab screening costs. Here we introduce AMPlify, an attentive deep learning model for AMP prediction, and demonstrate its utility in prioritizing peptide sequences derived from the *Rana [Lithobates] catesbeiana* (bullfrog) genome. We tested the bioactivity of our predicted peptides against a panel of bacterial species, including representatives from the World Health Organization’s “priority pathogens” list. Four of our novel AMPs were active against multiple species of bacteria, including a multi-drug resistant isolate of carbapenemase-producing *Escherichia coli*, demonstrating the utility of tools like AMPlify in our fight against antibiotic resistance.

## Introduction

As reported by the World Health Organization (WHO), the decreasing effectiveness of antibiotics and other antimicrobial agents indicates the world is at a risk of entering a “post-antibiotic” era^1^. To counter this threat, new drugs or effective substitutes for conventional antibiotics are urgently needed. Antimicrobial peptides (AMPs) are one such alternative. AMPs are host defense molecules produced by all forms of life, including multicellular organisms as part of their innate immunity against microbes. Within their respective hosts, AMPs have co-evolved with microorganisms to serve as a defense against bacterial^2^, fungal^3^ and even viral infections^4^. Unlike most conventional antibiotics, which have specific functional or structural targets, AMPs act directly on the biological membranes of microorganisms or modulate the host immunity to enhance defense against microorganisms^5^. Also, they act faster than conventional antibiotics, have a narrower active concentration window for killing, and do not typically damage the DNA of their targets^6^. As a result, they do not induce resistance to the extent that is observed with conventional antibiotics^7^. Nevertheless, if bacteria are exposed to AMPs for extended periods of time, they can and do develop resistance even to AMP-based drugs including the last resort and life-saving drug, Colistin^6^. Hence, fast and accurate methods would be valuable tools to discover and design effective AMPs to enhance our repertoire of alternative therapeutics.

Direct, large scale discovery of novel AMPs through wet lab screening is time-consuming, labor-intensive and costly^8^. For these reasons, various computational models have been developed over the last few years^8^ to streamline *in silico* AMP prediction. Despite the rapid progress in the field, currently available models still have substantial room for improvement.

The AMP prediction module in the Collection of Antimicrobial Peptides (CAMP) database^9^ includes four different models: random forest, support vector machine, discriminant analysis, and a single-hidden-layer feed-forward neural network with 64 designed features^10^. The iAMP-2L online server adopts fuzzy *K*-nearest neighbor algorithm, taking pseudo amino acid compositions (PseAAC) with five physicochemical properties as input features to predict AMPs as well as their potential microorganism targets^11^. The iAMPpred online server for AMP prediction and classification is based on support vector machine that uses PseAAC with compositional, physicochemical, and structural features^12^. All three of these models employ conventional machine learning methods and rely on pre-designed features, requiring prior expertise in AMP structure and mechanism for effective engineering.

Alternatively, deep learning methods can automatically learn high-level features and usually outperform conventional methods in many bioinformatics tasks^13^. For AMP prediction, Veltri and co-workers introduced a deep neural network classifier with embedding, convolutional, max pooling, and recurrent layers which is available as an online server, AMP Scanner Vr.2, as its user interface^14^. While AMP Scanner Vr.2 outperforms the conventional machine learning methods cited above, we note that the chosen neural network architecture faces limitations in extracting long-range information. Common deep learning methods for sequence classification include recurrent neural networks (RNNs) and convolutional neural networks (CNNs), as employed in combination by AMP Scanner Vr.2. RNNs are able to learn remote dependencies inside a sequence, but suffer from gradient vanishing problems^15^. Similarly, while CNNs can extract local information well, long-range dependencies are ignored^16^.

Recently, deep neural networks with attention mechanisms have gained momentum, notably for natural language processing^17–19^ and computer vision^20^ applications. Attention mechanisms, as the name suggests, are inspired by our brains’ ability to prioritize segments of information when processing textual or visual input. In sequence analysis, attention mechanisms are modeled by weights assigned to different positions in a sequence. These weights amplify or attenuate information from a given position to help encode the global information of the sequence.

Here, we introduce AMPlify, an attentive deep learning model that improves *in silico* AMP prediction by applying two types of attention mechanisms on top of a bidirectional long short-term memory^21–23^ (Bi-LSTM) layer (Fig. 1). The Bi-LSTM layer in the model, as a variant of RNN, encodes positional information from the input sequence in a recurrent manner. Subsequently, the multi-head scaled dot-product attention^18^ (MHSDPA) layer computes a refined representation of the sequence using multiple weight vectors. The last hidden layer of context attention^19^ (CA) generates a single summary vector using weighted average, learning contextual information gained from the previous layer. The AMPlify model is trained on a set of known AMPs and a select list of non-AMP sequences, and adopts ensemble learning to further improve its performance. To the best of our knowledge, AMPlify is the first machine learning application that applies attention mechanisms for *in silico* AMP prediction.

**Figure 1:**
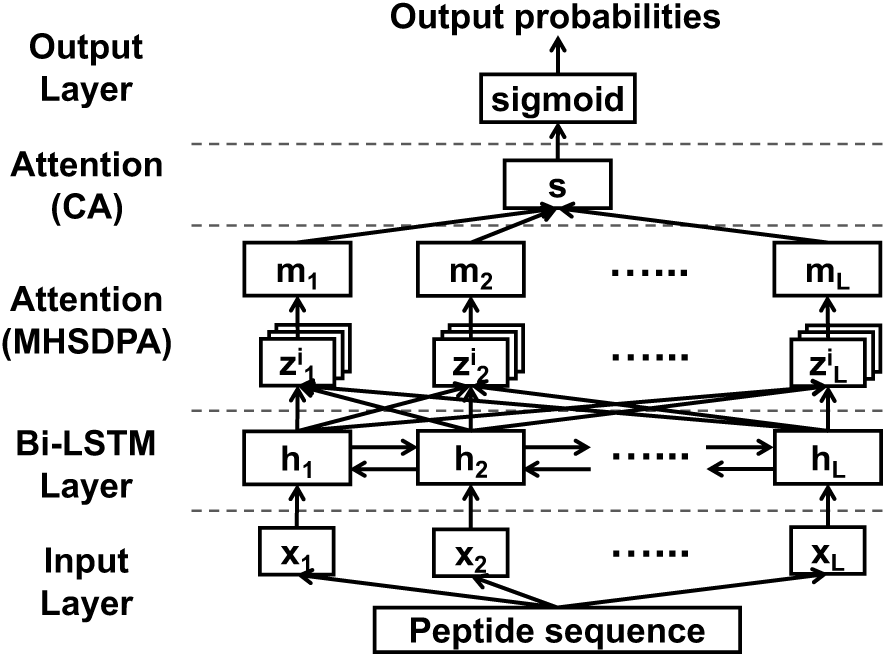
Model architecture of AMPlify. Residues of a peptide sequence are one-hot encoded and passed to three hidden layers in order: the bidirectional long short-term memory (Bi-LSTM) layer, the multi-head scaled dot-product attention (MHSDPA) layer and the context attention (CA) layer. The output layer generates the probability that the input sequence is an AMP.

To illustrate the utility of our model, a discovery pipeline based on AMPlify was used to mine the AMP-rich North American bullfrog (*Rana [Lithobates] catesbeiana*) genome for novel AMPs. In our tests, AMPlify outperformed state-of-the-art methods, and successfully identified novel AMPs with biological activity *in vitro*. The WHO has a published list of “priority pathogens” for which new antibiotics are urgently needed^24^. This list includes bacterial species that are increasingly resistant to multiple antibiotics. In the current study, we tested the efficacy of our discovered, putative AMPs against selected WHO-prioritized pathogens, including: (1) *Pseudomonas aeruginosa* and *Escherichia coli* strains, including a multi-drug resistant (MDR) carbapenemase-producing (CPO) strain of *E. coli* reflective of WHO’s “Priority 1” pathogens; and (2) a *Staphylococcus aureus* strain reflective of WHO’s “Priority 2” methicillin-resistant (MRSA) and vancomycin-resistant (VRSA) strains. A *Streptococcus pyogenes* strain was included as an additional Gram-positive bacterial species that causes human disease and has demonstrated antibiotic resistance^25^. Four of the 16 novel AMPs discovered show considerable antimicrobial potency against one or more of the organisms examined, including the clinical MDR isolate of CPO *E. coli.* These results highlight the potential of AMPlify to accelerate AMP discovery, the first step towards facilitating peptide-based therapeutics.

## Results

### Evaluation of model architecture

To demonstrate the effectiveness of each component within our model, we evaluated the model architecture starting from a single Bi-LSTM layer and then gradually adding attention layers over it. Supplementary Table 1 compares different model architectures using stratified 5-fold cross-validation on the training set with regard to five different measures of accuracy, sensitivity, specificity, F1 score and area under the receiver operating characteristic curve (AUROC). The first section of the table compares the performance of the complete architecture of AMPlify, with and without ensemble learning, with simpler variations which include fewer hidden layers. The architecture of the only deep learning based comparator, AMP Scanner Vr.2, was cross-validated on our training set for comparison using two different stopping settings: the optimal fixed number of epochs as stated in their manuscript^14^ and early stopping as described in this paper (Supplementary Table 1, second section). Although overall performance is not strongly influenced by early stopping, it does lead to smaller performance variability as measured by standard deviation values in tests, indicating that the model trained using early stopping is more robust than using a default of 10 epochs.

By adding a single CA layer atop the Bi-LSTM layer, the model performs similarly to AMP Scanner Vr.2 based on cross-validation results, with differences smaller than 1% in all metrics except specificity (<1.4%). After inserting an MHSDPA layer in the middle, the cross-validation results reach 91.70% in accuracy, 91.40% in sensitivity, 92.00% in specificity, 91.68% in F1 score, and 96.92% in AUROC – an overall improvement compared with the architecture without this layer. This indicates that the attention layer learns discriminating features of sequences processed by the Bi-LSTM layer. We note that the final AMPlify architecture already outperforms the AMP Scanner Vr.2 architecture in all metrics in our cross-validation tests. After applying ensemble learning to the proposed architecture, the cross-validation performance is further improved to 92.79% for accuracy, 92.12% for sensitivity, 93.47% for specificity, 92.74% for F1 score and 97.44% for AUROC.

Paired student t-tests based on our cross-validation results indicate statistically significant increase in performance by AMPlify over AMP Scanner Vr.2 (early stopped) with regard to all five metrics (p<0.05) except specificity (p=0.068). Similarly, the better performance of AMPlify without ensemble learning (i.e. Bi-LSTM+MHSDPA+CA) over the simple Bi-LSTM model is statistically significant in all metrics (p<0.05), suggesting that the attention layers play important roles in the model’s improvement.

The superior performance of AMPlify when cross-validated on the dataset provided by AMP Scanner Vr.2 further suggests that the deep neural network architecture chosen in AMPlify is better for the task of AMP prediction than the architecture of AMP Scanner Vr.2 (Supplementary Note 1, Supplementary Table 2).

### Comparison with state-of-the-art methods

With the set of hyperparameters tuned through stratified 5-fold cross-validation, the final model of AMPlify was trained using the entire training set, with each of the five sub-models trained on five different subsets. AMPlify, along with its sub-models, were compared on our test set with three other state-of-the-art tools: iAMP-2L^11^, iAMPpred^12^ and AMP Scanner Vr.2^14^ (Table 1). All the tools were evaluated with their original trained models reported. AMP Scanner Vr.2 could be trained using third party datasets through a utility provided by the authors (personal communication with Daniel Veltri), and was re-trained on our training set with two different stopping conditions, as previously stated.

**Table 1:**
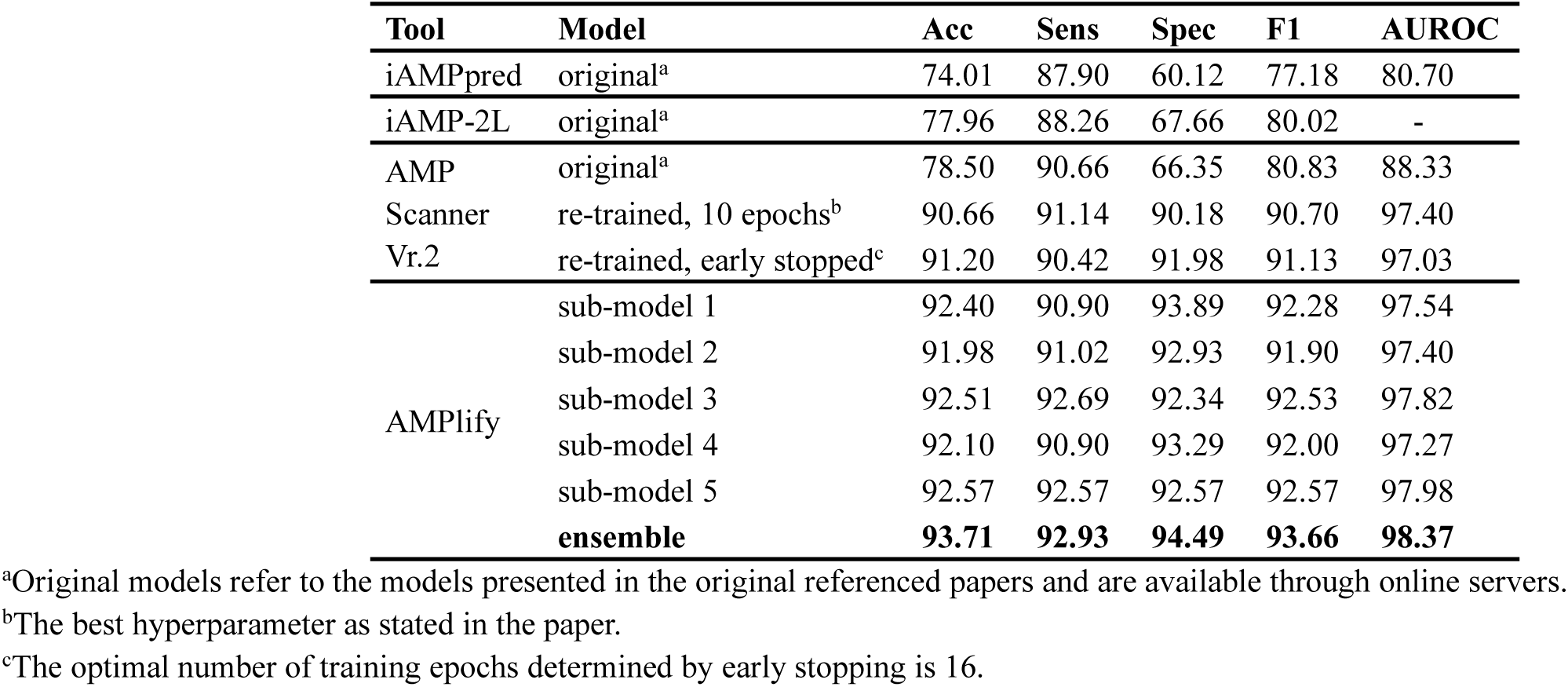
Performance comparison among different tools on the test set. Performance of different tools are presented with five metrics in percentage: accuracy (acc), sensitivity (sens), specificity (spec), F1 score (F1) and area under the receiver operating characteristic curve (AUROC).

Among the original models of the three comparators, AMP Scanner Vr.2 performs the best on our data in general, except that its specificity is 1.31% lower than iAMP-2L. The accuracy, specificity, F1 score, and AUROC of AMP Scanner Vr.2 were all improved after re-training, with only small changes in sensitivity (<0.5%). However, results show that AMPlify outperforms all the competing models, including the two “re-trained” versions of AMP Scanner Vr.2. AMPlify achieves the highest accuracy (93.71%), F1 score (93.66%) and AUROC (98.37%), which beats those “re-trained” versions of AMP Scanner Vr.2 by 2.51%, 2.53% and 0.97% respectively. AMPlify also shows the highest sensitivity (92.93%) and specificity (94.49%), suggesting that the model can concurrently reduce false negative and false positive predictions.

Further, all five sub-models of AMPlify yield favourable performance in accuracy (91.98%–92.57%), specificity (92.34%–93.89%) and F1 score (91.90%–92.57%), despite each sub-model being trained on 80% of the entire training set (see **Methods**). The sensitivity values of the five sub-models range from 90.90% to 92.69%, with two of them being better than the performance of all comparators, while the remaining three being slightly lower than the performance of the “re-trained, 10 epochs” model of AMP Scanner Vr.2 (<0.25%). Still, the lower standard deviation values from cross-validation analysis indicate that those single sub-models of AMPlify are more robust compared with the “re-trained, 10 epochs” model of AMP Scanner Vr.2 (Supplementary Table 1). Similarly, our sub-models score higher than the comparators in AUROC, except one of them being on par with the “re-trained, 10 epochs” version of AMP Scanner Vr.2 and another scoring lower by 0.13%. The specificity values of the original models of the three comparators are relatively low (<70%), likely due to their less stringent selection criteria when building their non-AMP sets. The specificity values of AMP Scanner Vr.2 improved substantially after being re-trained on our strictly selected training set (90.18% and 91.98% depending on the number of epochs trained, Table 1), although they are still lower than those of all the sub-models and the final ensemble model of AMPlify. We have also conducted a cross-comparison of AMPlify with AMP Scanner Vr.2, re-training and testing our tool on the dataset provided by the AMP Scanner Vr.2 publication^14^, which illustrated the improved learning capability of our chosen architecture for the task of AMP prediction (Supplementary Note 1, Supplementary Table 3, Supplementary Fig. 1).

In order to compare the classification performance of each tool with regard to different classification thresholds, Fig. 2a presents a series of receiver operating characteristic (ROC) curves for the models compared. The AUROC results shown in Table 1 correspond to these ROC curves. Note that the iAMP-2L online server does not allow for parameterization, hence the tool is represented by a single data point. The ROC curves indicate that AMPlify is Pareto-optimal in our tests for any classification threshold.

**Figure 2:**
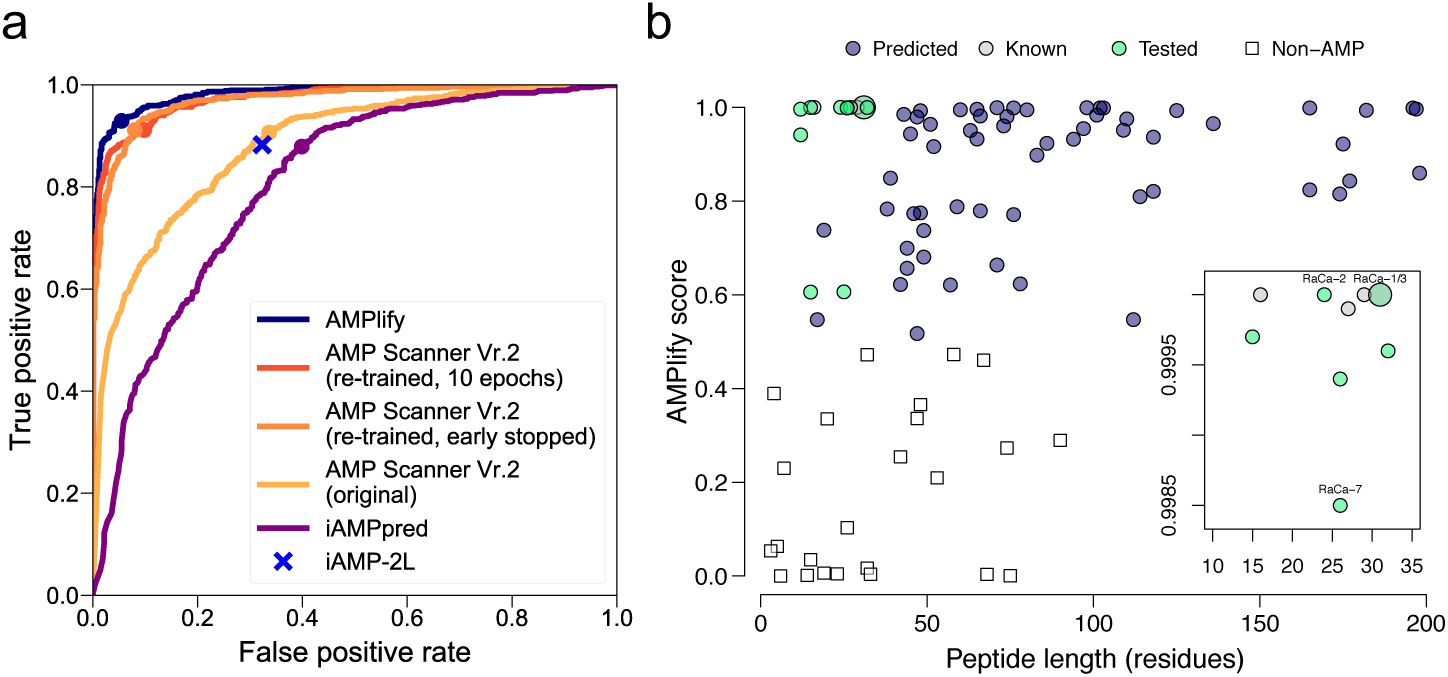
Visualization of AMPlify model performance and the AMP discovery pipeline application results. **(a)** Receiver operating characteristic (ROC) curves of AMPlify and competing models are plotted, with round dots marking the performance at the threshold of 0.5. The iAMP-2L online server only output labels of AMP/non-AMP without the corresponding probabilities, so it appears as a single point on the plot. **(b)** AMPlify prediction scores against peptide lengths of 101 sequences analyzed by AMPlify. Inset shows amplified view of the upper left region of the plot to enhance visualization of the majority of the selected sequences.

### AMP discovery

Previous studies have shown that the skin secretions of amphibians are rich in AMPs, which help prevent infection by harmful microorganisms^26^. For this reason, mining the genomes of various frog species for novel AMPs is an attractive proposition. To demonstrate AMPlify’s practical application, it was embedded into a bioinformatics AMP discovery pipeline to find novel AMPs from the North American bullfrog (*Rana [Lithobates] catesbeiana*) genome^27,28^. For antimicrobial susceptibility testing (AST), we focus on cationic AMPs acting directly on biological membranes, the activities of which can be directly observed *in vitro*. Most amphibian AMP precursors possess highly conserved N-terminal prepro regions and hypervariable C-terminal antimicrobial domains^26^. Based on this, we identified candidate precursors from the bullfrog genome using homology search and genome annotation tools. We then derived candidate mature sequences from those precursors to use as input for AMPlify. Please refer to the **Methods** section for pipeline details. This resulted in 101 candidate mature sequences, which we fed into AMPlify, predicting 75 of them to be putative AMPs. We selected peptides between five to 35 amino acids in length with a positive charge for further analysis, yielding a final list of 16 peptides (Table 2), five of which were previously reported sequences from literature or AMP databases^28–30^. The remaining 11 peptides were synthesized and evaluated *in vitro*. Fig. 2b shows a visualization of AMPlify prediction results for the 101 candidate mature sequences.

**Table 2:**
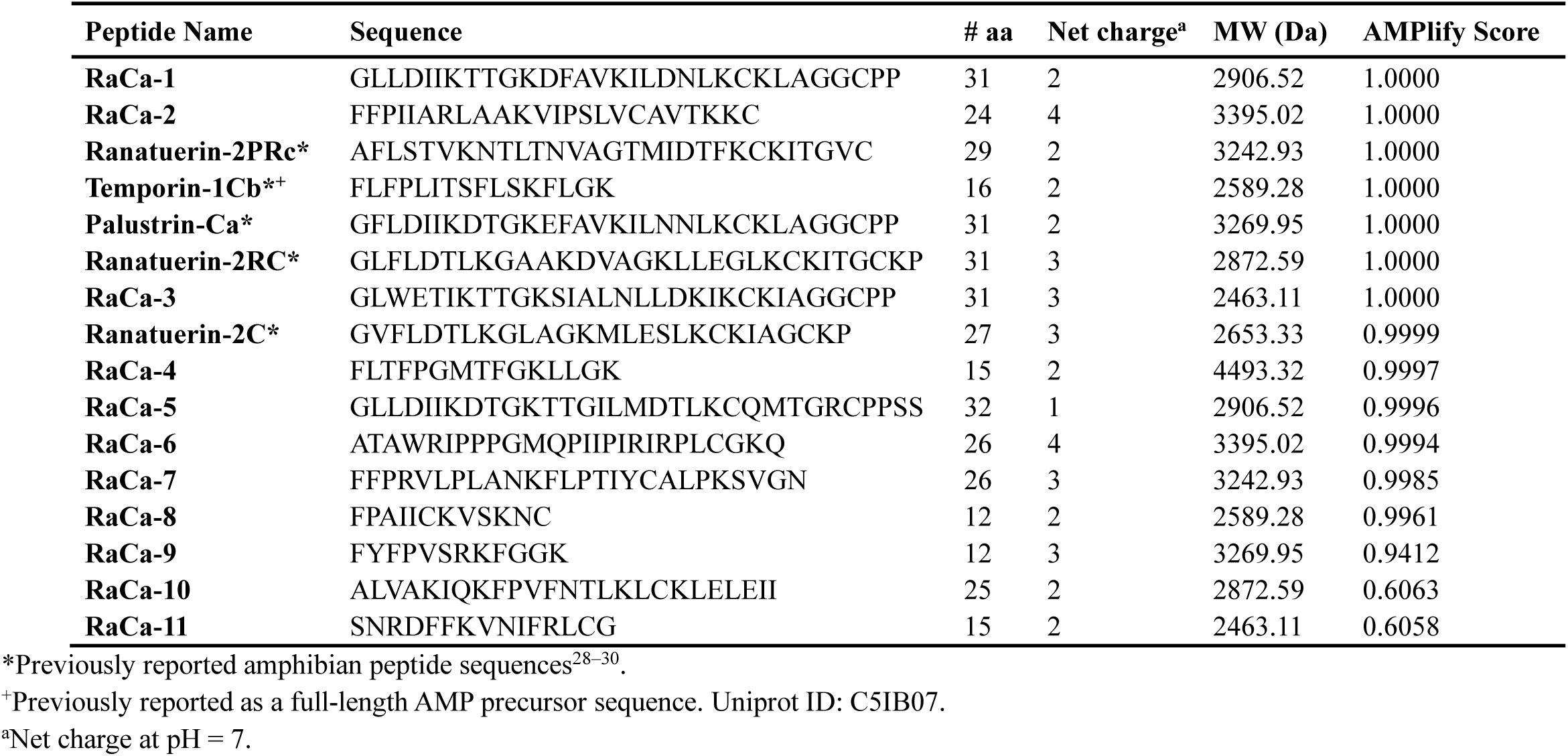
Putative and reported AMP sequences discovered from *Rana [Lithobates] catesbeiana*. Genomic and transcriptomic resources from *Rana [Lithobates] catesbeiana*^27^ were mined using the AMP discovery pipeline based on AMPlify. Top-scoring peptide sequences were selected for synthesis and validation *in vitro*.

### Antimicrobial susceptibility testing (AST)

A panel comprised of six bacteria was selected to test candidate AMP sequences identified using AMPlify: *Staphylococcus aureus* ATCC 6538P, *Streptococcus pyogenes* (unknown strain; hospital isolate), *Pseudomonas aeruginosa* ATCC 10148, *Escherichia coli* ATCC 9723H and ATCC 29522, and a multi-drug resistant (MDR) carbapenemase-producing New-Delhi metallobetalactamase (CPO-NDM) *Escherichia coli* clinical isolate. *E. coli* ATCC 29522 was used as a “wild-type” drug susceptible control strain. Results from AST are presented in Table 3.

**Table 3:**
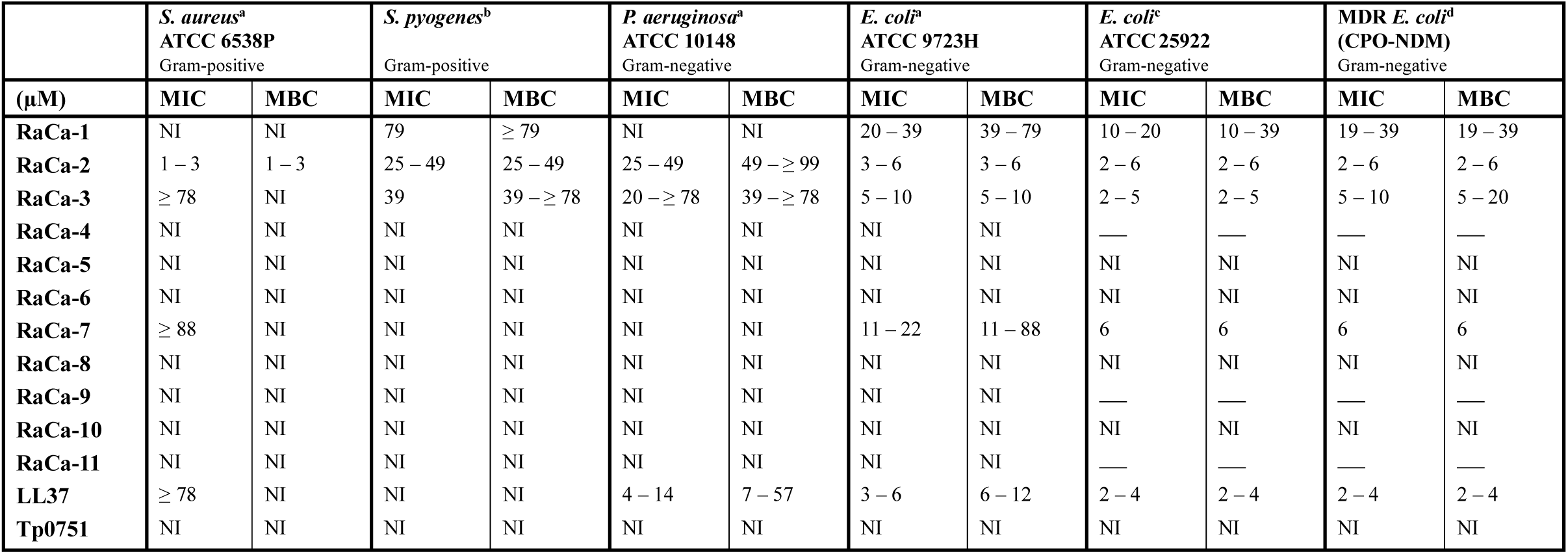
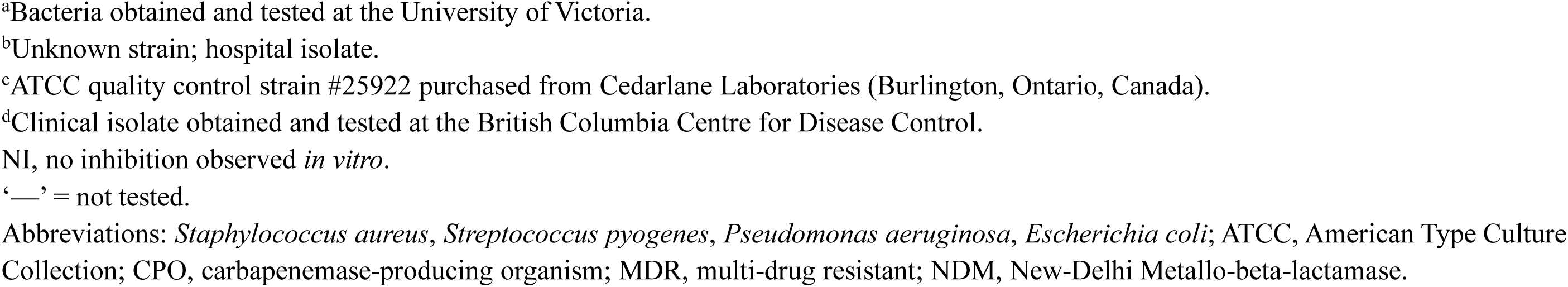
Minimum inhibitory concentrations (MIC) and minimum bactericidal concentrations (MBC) of selected AMP candidates following antimicrobial susceptibility testing (AST) *in vitro*. Candidate antimicrobial peptides were synthesized and purchased from Genscript. AST, and MIC/MBC determination was performed as outlined by the Clinical and Laboratory Standards Institute (CLSI)^47^, with modification as recommended by R.E.W. Hancock^48^. Data is presented as the lowest effective peptide concentration range (µM) observed in three independent experiments. LL37, human cathelicidin and a peptide from Tp0751 from *T. pallidum* were used as the positive and negative control peptides^28^, respectively.

|  | *S. aureus*<sup>a</sup><br>ATCC 6538P<br>Gram-positive |  | *S. pyogenes*<sup>b</sup><br>Gram-positive |  | *P. aeruginosa*<sup>a</sup><br>ATCC 10148<br>Gram-negative |  | *E. coli*<sup>a</sup><br>ATCC 9723H<br>Gram-negative |  | *E. coli*<sup>c</sup><br>ATCC 25922<br>Gram-negative |  | MDR *E. coli*<sup>d</sup><br>(CPO-NDM)<br>Gram-negative |  |
| --- | --- | --- | --- | --- | --- | --- | --- | --- | --- | --- | --- | --- |
| (μM) | MIC | MBC | MIC | MBC | MIC | MBC | MIC | MBC | MIC | MBC | MIC | MBC |
| **RaCa-1** | NI | NI | 79 | ≥ 79 | NI | NI | 20 – 39 | 39 – 79 | 10 – 20 | 10 – 39 | 19 – 39 | 19 – 39 |
| **RaCa-2** | 1 – 3 | 1 – 3 | 25 – 49 | 25 – 49 | 25 – 49 | 49 – ≥ 99 | 3 – 6 | 3 – 6 | 2 – 6 | 2 – 6 | 2 – 6 | 2 – 6 |
| **RaCa-3** | ≥ 78 | NI | 39 | 39 – ≥ 78 | 20 – ≥ 78 | 39 – ≥ 78 | 5 – 10 | 5 – 10 | 2 – 5 | 2 – 5 | 5 – 10 | 5 – 20 |
| **RaCa-4** | NI | NI | NI | NI | NI | NI | NI | NI | — | — | — | — |
| **RaCa-5** | NI | NI | NI | NI | NI | NI | NI | NI | NI | NI | NI | NI |
| **RaCa-6** | NI | NI | NI | NI | NI | NI | NI | NI | NI | NI | NI | NI |
| **RaCa-7** | ≥ 88 | NI | NI | NI | NI | NI | 11 – 22 | 11 – 88 | 6 | 6 | 6 | 6 |
| **RaCa-8** | NI | NI | NI | NI | NI | NI | NI | NI | NI | NI | NI | NI |
| **RaCa-9** | NI | NI | NI | NI | NI | NI | NI | NI | — | — | — | — |
| **RaCa-10** | NI | NI | NI | NI | NI | NI | NI | NI | NI | NI | NI | NI |
| **RaCa-11** | NI | NI | NI | NI | NI | NI | NI | NI | — | — | — | — |
| **LL37** | ≥ 78 | NI | NI | NI | 4 – 14 | 7 – 57 | 3 – 6 | 6 – 12 | 2 – 4 | 2 – 4 | 2 – 4 | 2 – 4 |
| **Tp0751** | NI | NI | NI | NI | NI | NI | NI | NI | NI | NI | NI | NI |

<sup>a</sup>Bacteria obtained and tested at the University of Victoria.
<sup>b</sup>Unknown strain; hospital isolate.
<sup>c</sup>ATCC quality control strain #25922 purchased from Cedarlane Laboratories (Burlington, Ontario, Canada).
<sup>d</sup>Clinical isolate obtained and tested at the British Columbia Centre for Disease Control.
NI, no inhibition observed *in vitro*.
‘—’ = not tested.
Abbreviations: *Staphylococcus aureus*, *Streptococcus pyogenes*, *Pseudomonas aeruginosa*, *Escherichia coli*; ATCC, American Type Culture Collection; CPO, carbapenemase-producing organism; MDR, multi-drug resistant; NDM, New-Delhi Metallo-beta-lactamase.

The 11 putative AMP sequences were selected for *in vitro* AST experiments, and four of them had antimicrobial activity against the targets tested: RaCa-1, RaCa-2, RaCa-3, and RaCa-7. RaCa-1 was antibacterial against all *E. coli* strains tested (MIC = 10–39 µM, MBC = 10–79 µM). RaCa-1 also showed minimal antimicrobial activity against *S. pyogenes* (MIC/MBC ≥ 79 µM) with no observed inhibition against the *S. aureus* and *P. aeruginosa* isolates. RaCa-2 and RaCa-3 inhibited all bacterial strains tested. RaCa-2 possessed the strongest antibacterial activity against *S. aureus* and *E. coli* isolates, preventing growth of both species of bacteria at concentrations of 1–3 µM and 2–6 µM, respectively. Specifically, this peptide was bactericidal against *E. coli* ATCC 9723H (MIC/MBC = 3–6 µM), with similar activity observed against *E. coli* ATCC 25922 and the MDR *E. coli* CPO-NDM isolates (MIC/MBC = 2–6 µM). RaCa-2 was also the only AMP tested to have robust bactericidal action against both of the tested Gram-positive bacteria, *S. aureus* (MIC/MBC = 1–3 µM) and *S. pyogenes* (MIC/MBC = 25–49 µM). Comparably, RaCa-3 was considerably potent *in vitro* against *S. pyogenes* (MIC = 39 µM, MBC = 39–≥78 µM), *P. aeruginosa* (MIC = 20– ≥78 µM, MBC = 39–≥78 µM), *E. coli* (MIC = 2–10 µM, MBC = 2–20 µM), and to a lesser extent *S. aureus* (MIC ≥ 78 µM, MBC = NI). RaCa-7 was active against all strains of *E. coli* (MIC = 6–22 µM, MBC = 6–88 µM), with minimal inhibition of *S. aureus* (MIC ≥ 88 µM, MBC = NI), and no activity against the other two species. Overall, the four novel AMP sequences displayed the strongest activity against the tested *E. coli* strains. RaCa-2, RaCa-3, and RaCa-7 all had significant antibacterial action against the MDR *E. coli* (CPO-NDM), each inhibiting bacterial growth at ≤ 10 µM. Of particular note, there was little or no observed shift in MIC and MBC values when comparing the CPO-NDM *E. coli* isolate to the ATCC 25922 “wild-type” control strain.

The positive control peptide LL37^28^ displayed potent antimicrobial activity against all strains of *E. coli* (MIC = 2–6 µM, MBC = 2–12 µM) and *P. aeruginosa* (MIC = 4–14 µM, MBC = 7–57 µM). However, this peptide had minimal activity against the tested strains of *S. aureus* and *S. pyogenes*. The negative control peptide, Tp0751, a non-functional truncated section of a *Treponema pallidum* protein with similar characteristics to AMPs^31^, was inactive against all organisms.

## Discussion

Here we present AMPlify, a robust attentive deep learning model for AMP prediction, and demonstrate its utility in identifying novel AMPs with broad antimicrobial activities. To the best of our knowledge, AMPlify is the first method to apply attention mechanisms inside a deep neural network for this task. It implements ensemble learning by partitioning its training set – a novel approach – and outperforms existing machine learning methods, including a leading deep learning-based model. The two attention mechanisms are inspired by how humans perceive natural language, paying closer attention to regions or words of interest in a sentence. We have observed that sub-models of AMPlify were able to outperform the state-of-the-art methods without ensemble learning, and trace the source of this favourable performance to the inclusion of attention layers.

Although machine learning methods in general, and AMPlify in particular, perform well in predicting AMPs, their performance can be limited by a paucity of detailed AMP sequence data available for training. First, the models do not usually consider the potential target microorganisms for the predicted AMPs. Although some methods report success at that level of granularity using public data^11,12^, incomplete and incorrect annotations in AMP databases are confounding. Second, the models cannot distinguish whether an AMP acts directly on biological membranes and/or by modulating the host immunity, since there is no consistently available data on these features. AMPs acting only in the latter mode require separate assays and might differ in activity within different species. Third, the size of the training data is still small relative to the data typically employed in most deep learning applications. We expect this limitation to be gradually alleviated as more AMPs are discovered. Although the size of the training data is unlikely to match what is available in natural language processing, image classification, and social network analysis domains, to name a few, AMP prediction tools can still find practical applications as demonstrated here.

Using AMPlify, four novel AMPs were identified with proven activity against a variety of bacterial isolates. Promisingly, three of the four presented AMPs demonstrate antibacterial activity against the MDR *E. coli* tested, and there was little or no observed shift in MIC when comparing the MDR and drug-susceptible strains. This suggests that the mechanism-of-action of these AMPs is unlike those used by conventional antibiotics. Thus, the AMPs presented in the current study have the potential to be used in future drug and clinical development studies as peptide-based substitutes to classical antibiotics. Although several candidates identified using this pipeline did not show any *in vitro* activity against the bacteria tested, they still may possess activity against other bacterial species or other microorganisms (e.g. fungi, virus), or may demonstrate activity *in vivo* via host immune response modulation.

Of course, the utility of AMPlify is not limited to discovering AMPs from the bullfrog genome, and may also be used to interrogate additional genomes. AMPlify can be generically applied to any input sequence. As such, it has the potential to play a role in *de novo* AMP design or enhancement. In conclusion, with their various use cases, we foresee tools like AMPlify as being instrumental in expanding the current arsenal of antimicrobial agents effective against WHO “priority pathogens”.

## Methods

### Generation of the datasets

We used publicly available AMP sequences to train and test AMP predictors. In order to build a non-redundant AMP dataset, we first downloaded all available sequences from two manually curated databases: Antimicrobial Peptide Database^32^ (APD3, http://aps.unmc.edu/AP) and Database of Anuran Defense Peptides^30^ (DADP, http://split4.pmfst.hr/dadp). Since APD3 is being frequently updated, we used a static version that was scraped from the website on March 20, 2019 comprising 3,061 sequences. Version 1.6 of DADP contains 1,923 distinct mature AMPs. We concatenated these two sets and removed duplicate sequences, producing a non-redundant (positive) set of 4,173 distinct, mature AMP sequences, all 200 amino acid residues in length or shorter.

Training and testing binary classification models require a negative set, a collection of peptides known not to have any antimicrobial activity. Since there are no sequence catalogs for peptides devoid of antimicrobial activity, studies in the field typically select their non-AMP sequences from UniProt^33^ (https://www.uniprot.org). This may involve excluding several simple keywords (e.g. antimicrobial, antibiotic) to filter out potential AMPs^10,11^, or additionally removing all secretory proteins^14^ as AMPs are characteristically secreted peptides^34^. The former proposition is not sufficiently rigorous, because AMP annotation is not consistent and varies between sources. While the former approach may leave in the set some differently annotated AMPs, the latter creates a learning gap for the model regarding secretory proteins without antimicrobial activities. Thus, it is important to balance these two strategies when selecting non-AMP sequences.

We designed a rigorous selection strategy for our non-AMP sequences (Supplementary Fig. 2), using sequences from the UniProtKB/Swiss-Prot database^33^ (2019_02 release), which only contains manually annotated and reviewed records from the UniProt database. First, we downloaded sequences that are 200 amino acid residues or shorter in length, excluding those with annotations containing any of the 16 following keywords related to antimicrobial activities: {antimicrobial, antibiotic, antibacterial, antiviral, antifungal, antimalarial, antiparasitic, anti-protist, anticancer, defense, defensin, cathelicidin, histatin, bacteriocin, microbicidal, fungicide}. Second, duplicates and sequences with residues other than the 20 standard amino acids were removed. Third, a set of “potential AMP sequences” annotated with any of the 16 selected keywords were downloaded and compared with our candidate negative set. We noted instances where a sequence with multiple functions was annotated separately in multiple records within the database, and removed sequences in common between candidate non-AMPs and potential AMPs. The candidate non-AMP sequences were also checked against the positive set to remove AMP sequences that lack the annotation in UniProtKB/Swiss-Prot. Finally, 4,173 sequences were sampled from the remaining non-AMP set, matching the number and length distribution of sequences in the positive set. An exception to the length distribution matching occurred when the length of a particular AMP sequence did not have a perfect match in the set of 128,445 non-AMP sequences. In these instances, we chose the non-AMP sequence with the closest length. The matched length distributions were selected so that the model did not learn to distinguish classes based on sequence lengths.

The positive and negative sets were both split 80%/20% (3,338/835) into training and test sets, respectively. We note that AMP sequences in our test partition have no overlap with the training sets of iAMPpred and iAMP-2L, but do share 196 sequences with the training set of AMP Scanner Vr.2.

### Model implementation

AMPlify is implemented in Python 3.6.7, using Keras library 2.2.4^35^ with Tensorflow 1.12.0^36^ as the backend. The raw output of the model can be interpreted as a probability score, thus sequences with scores > 0.5 are considered as AMPs and those ≤ 0.5 as non-AMPs. We used binary cross-entropy as the loss function, and the “Adam”^37^ algorithm for optimizing weights. Dropout technique^38^ was applied during training to prevent the model from over-fitting. The original positive and negative training sets were both split into sets of {667, 667, 668, 668, 668} sequences, and five training and validation set pairs were constructed by leaving one set out for validation to build five sub-models. To optimize computational time and avoid overfitting, we applied early stopping during the training of each sub-model. The validation accuracy was monitored at each training epoch, and the training process was stopped if there appeared to be no improvement for the next 50 epochs. The model weights from the epoch with the best validation accuracy were selected as the optimal weights. The output probabilities from the five sub-models were averaged to yield an ensemble model.

Reflecting the composition of the sequences in the positive and negative sets, AMPlify only considers sequence lengths of 200 or shorter containing the 20 standard amino acid residues. The source code for AMPlify and the trained models are available at https://github.com/bcgsc/AMPlify.

### Hyperparameter tuning and model architecture

In deep neural networks, the optimal hyperparameters, unlike model weights, cannot be learned directly from the training process, but they do play an important role in model performance. Thus, various combinations of hyperparameters must be compared in order to select the best set. Here we applied stratified 5-fold cross-validation on the entire training set to tune the model and find the best set of hyperparameters for the model architecture, as well as for training settings, including dropout rates and optimizer settings. For a fair comparison, we kept the same splits of sequences within cross-validation across all hyperparameter combinations. During hyperparameter tuning, we monitored the average performance on the validation partitions of cross-validation. Note that these validation partitions within cross-validation are different from the validation sets for early stopping, while the latter being additionally derived from the training partitions during the cross-validation process. The set of hyperparameters with the highest average cross-validation accuracy was chosen to train the final prediction model.

The AMPlify architecture includes three main components: (1) a bidirectional long short-term memory (Bi-LSTM) layer, (2) a multi-head scaled dot-product attention (MHSDPA) layer, and (3) a context attention (CA) layer (Fig. 1). To convert the original peptides into a mathematically processable format, each sequence is represented by a series of one-hot encoded vectors over an alphabet of 20 amino acids, yielding **x**_**1**_, **x**_**2**_, …, **x**_**L**_, where *L* is the length of the sequence and each **x**_**t**_ is a 20-dimensional vector of zeros and ones with ∥**x**_**t**_∥_**1**_ = 1 (*t* = 1,2, …, *L*). The Bi-LSTM layer takes those one-hot encoded vectors as input and encodes positional information for each residue from both forward and backward directions, and the output vector for each residue is represented as a concatenation of the vectors from both directions. The best tuned dimensionality for each direction of Bi-LSTM layer was 512, resulting in the entire Bi-LSTM layer to be *d*_*h*_ = 512 × 2 = 1024 dimensional. Outputs from all timesteps of the Bi-LSTM layer are returned as the input for the next layer. The best tuned dropout rate of 0.5 was applied to the input of the Bi-LSTM layer. Encoding from the Bi-LSTM layer for residues within a given sequence can be represented as a series of vectors 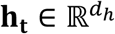 (*t* = 1,2, …, *L*), and the entire sequence can be represented as a matrix with all **h**_**t**_s packed as

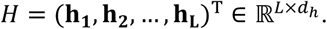

Next, the MHSDPA layer searches for relations between different residues in *n* different representation subspaces^18^ (i.e. different attention heads) to further encode the sequence, where *n* is a hyperparameter to be tuned. Each residue first gets an intermediate representation within each head by calculating a weighted average over transformed vectors of all residues derived from their Bi-LSTM representations. The results from each head are then concatenated and mapped back to the original dimensionality. We adopted Vaswani and co-workers’ approach^18^ to calculate the attention weights and outputs for the MHSDPA layer. Different from their approach, we added ReLU activation functions and bias terms to all linear transformations, which yielded better performance in our AMP prediction task. The implementation was adapted from the GitHub repository at https://github.com/CyberZHG/keras-multi-head.

In further detail, to obtain attention weights for different residues of a sequence within a head *i*, we calculate a set of queries 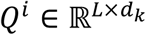, keys 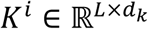, and values 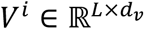 by transforming *H* as follows:

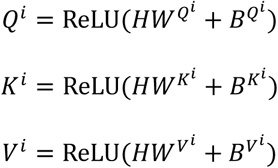

where 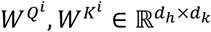 and 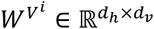 are weight matrices, and 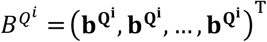, 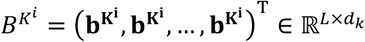 and 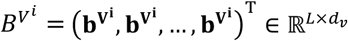 are bias matrices. We set transformation dimensions as *d*_*k*_ = *d*_*v*_ = *d*_*h*_/*n* following what has been previously proposed^18^. A square matrix *A*^*i*^ ∈ ℝ^*L*×*L*^, which contains weight vectors to calculate intermediate representations of all residues within head *i*, is computed as:

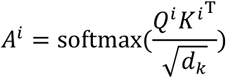

where dot-product of queries and keys are scaled by a factor 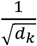, and the softmax function is applied to each row of the matrix for normalization. The intermediate representation of the sequence within head *i* is then computed by:

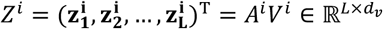

where each single vector 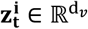 (*t* = 1,2, …, *L*) denotes the intermediate representation of each residue of the sequence with dimensionality *d*_*v*_. The concatenated matrix 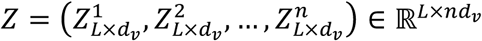 is further transformed to get the final output of the current layer as follows:

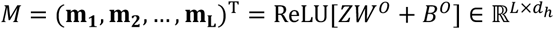

where 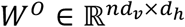 is a weight matrix and 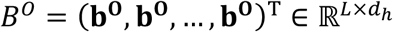 is a bias matrix. Each vector 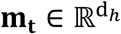 (*t* = 1,2, …, *L*) denotes the new representation of each residue of the sequence with dimensionality *d*_*h*_. The best head number tuned for this layer was *n* = 32, with *d*_*k*_ = *d*_*v*_ = 32 accordingly.

Finally, the CA layer gathers information from the MHSDPA layer by weighted averaging *L* encoded vectors into a single summary vector **s**. We followed Yang and co-workers’ approach^19^, and the implementation was adapted from the GitHub repository at https://github.com/lzfelix/keras_attention. The weight vector **α** ∈ ℝ^*L*^ can be obtained as:

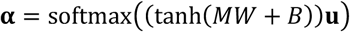

where 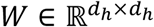 is a weight matrix, 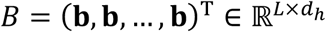 is a bias matrix, and 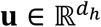 is a context vector, and the softmax function provides weight normalization. The summary vector 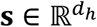 is then computed as:

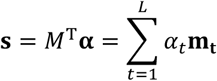

where *α*_*t*_ denotes each component in the weight vector. The summary vector summarizes information of the entire sequence into a single vector, and it is passed through the output layer with a sigmoid activation function for classification. The best tuned dropout rate of 0.2 was applied to the input of the CA layer during training.

In addition to the hyperparameters of the model architecture, the hyperparameters of the optimizer were optimized through cross-validation. The default setting with a learning rate of 0.001 in Keras was found to be the best for the AMP prediction task.

### Model evaluation

The performance of AMPlify was evaluated using five metrics: accuracy, sensitivity, specificity, F1 score and area under the receiver operating characteristic curve (AUROC).

The architecture of AMPlify was compared with its simpler variations with fewer hidden layers using stratified 5-fold cross-validation on the training set to measure the value added by each layer as the architecture grew more complex. The final version of AMPlify trained on the entire training set, as well as its five sub-models, were compared with three other tools: iAMP-2L^11^, iAMPpred^12^ and AMP Scanner Vr.2^14^, on the test set we built. All comparators were evaluated with their original models online.

In addition, as the only comparator with methods for re-training, AMP Scanner Vr.2 was cross-validated and re-trained on our training set for a more fair comparison. Since our dataset is slightly different from those used by other methods, the number of epochs required to get a deep learning model well trained on different datasets might differ. Keeping all other hyperparameters the same as the original model, we cross-validated and re-trained AMP Scanner Vr.2 with two different stopping settings: using the optimal fixed number of epochs as reported^14^, and using early stopping.

### AMP discovery pipeline

A primarily homology-based approach was used to supply novel candidate AMP sequences to AMPlify for further evaluation. The pipeline and its results are summarized in Supplementary Fig. 3 and are detailed below.

Sequences matching the search phrase “((antimicrobial) AND precursor) AND amphibian” were downloaded from the NCBI Nucleotide database on January 4th, 2016 and aligned to the draft bullfrog genome^27^ (version 3) using GMAP^39^ (version 20170424) with the following parameters: -A --max-intronlength-ends=200000 -O - n20 --nofails.

To refine the putative AMP loci, the gene prediction pipeline MAKER2^40^ (version 2.31.8 running under PERL version 5.24.0 with augustus^41^ version 3.2.1, exonerate^42^ version 2.2.0, genemark^43^ version 2.3c, and snap^44^ version 2006-07-28) was applied to the 231 genomic scaffolds with alignment hits from GMAP using default settings. The MAKER2 pipeline can use orthogonal evidence from related protein or transcript sequences when available to generate a list of high confidence genes. Protein evidence consisted of three sets of sequences: sequences matching the search phrase “((antimicrobial) AND precursor) AND amphibian” from the NCBI protein database that were downloaded on December 31st, 2015; experimentally validated non-synthetic amphibian antibacterial peptide sequences downloaded from CAMP^9^ on March 4th, 2016; and sequences from APD3^32^ downloaded on September 29th, 2017. For transcript evidence, the set of cDNA sequences supplied to GMAP above was supplemented with selected bullfrog transcript sequences from the Bullfrog Annotation Reference for the Transcriptome^27^ (BART). Blastn^45^ (version 2.31.1) was used to align the initial cDNA sequences from NCBI to BART, and BART sequences with an alignment of greater than 90% identity and 100% coverage were selected. A custom repeat element library was constructed from predicted repeats previously identified in the bullfrog genome^27^ and supplied to MAKER2 for use by RepeatMasker^46^. The annotation pipeline was run with the snap hidden Markov model that produced the version 2 bullfrog gene predictions^27^.

The MAKER2 gene predictions were filtered in two stages. First, sequences containing the highly conserved lysine-arginine enzymatic cleavage motif were selected and the sequence of the putative mature peptide, produced via *in silico* cleavage at the C-terminal side of the cleavage motif, was extracted. Second, only putative mature sequences of 200 amino acid residues or less were included. Sequences with non-standard amino acid residues were excluded. The resulting peptide sequences from these filters were fed into AMPlify for prediction. From the predicted putative AMPs, only short cationic sequences with lengths between five and 35 amino acid residues were chosen for synthesis and validation *in vitro*.

### Antimicrobial susceptibility testing (AST)

The 11 novel candidate AMP sequences predicted by AMPlify were selected for validation *in vitro*. Minimum inhibitory concentrations (MIC) and minimum bactericidal concentrations (MBC) were obtained using the AST procedures outlined by the Clinical and Laboratory Standards Institute (CLSI)^47^, with the recommended adaptations for testing cationic AMPs described by R.E.W. Hancock^48^. We prioritized short cationic sequences as shorter sequences are more structurally stable in various environments (e.g. *in vitro* and *in vivo*)^49^ lending to better therapeutic applicability.

#### Bacterial isolates

A panel of two Gram-positive and four Gram-negative bacterial isolates was generated to test predicted AMPs. *Staphylococcus aureus* ATCC 6538P, *Streptococcus pyogenes* (hospital isolate, unknown strain), *Pseudomonas aeruginosa* ATCC 10148, and *Escherichia coli* ATCC 9723H were obtained and tested at the University of Victoria. Additionally, a multi-drug resistant (MDR), carbapenemase-producing New-Delhi metallobetalactamase (CPO-NDM) clinical isolate of *Escherichia coli* was obtained from the BC Centre for Disease Control. *E. coli* ATCC 29522 was purchased from Cedarlane Laboratories (Burlington, Ontario, Canada) for comparison of AMP activity between a “wild type”, drug-susceptible control and the MDR strain. The latter two strains were tested at the BC Centre for Disease Control using identical AST procedures.

#### Determination of MIC

Bacteria were streaked onto nonselective nutrient agar from frozen stocks and incubated for 18–24 hours at 37°C. To prepare a standardized bacterial inoculum, isolated colonies were suspended in Mueller-Hinton Broth (MHB) and adjusted to an optical density of 0.08–0.1 at 600 nm, equivalent to a 0.5 McFarland standard and representing approximately 1–2 × 10^8^ CFU/mL. The inoculum was diluted 1/250 in MHB to the target concentration of (5 ± 3) × 10^5^ CFU/mL. Total viability counts from the final inoculum were examined to confirm the target bacterial density was obtained.

Selected candidate AMPs were purchased from Genscript (Piscataway, NJ), where they were synthesized using the vendor’s Flexpeptide platform. Lyophilized peptides were suspended in sterile ultrapure water or filter-sterilized 0.2% acetic acid as recommended by solubility testing reports provided with the GenScript synthesis. AMPs were diluted from 2560 to 5 µg/mL by a two-fold serial dilution in a 96-well polypropylene microtitre plate before 100 µl of the standardized bacterial inoculum of (5 ± 3) × 10^5^ CFU/mL was added to each well. This generated a final test range of 256 to 0.5 µg/mL. MIC values were reported as the peptide concentration that produced no visible bacterial growth after a 16–24 hour incubation at 37°C.

#### Determination of MBC

For each AMP dilution series the contents of the MIC well and the two adjacent wells containing two- and four-fold MIC were plated onto nonselective nutrient agar and incubated for 24 hours at 37°C. The concentration which killed 99.9% of the initial inoculum was determined to be the MBC.

## Acknowledgements

We would like to thank Dr. Hong Yu and Dr. Karuna Karunakaran for their generous efforts when establishing the laboratory at the BCCDC. We would also like to thank Dr. Anat Yanai for useful suggestions for the manuscript development, and Zhuyi Xue for helpful discussions on the design of the model. This work was supported by Genome BC and Genome Canada [281ANV]; and the National Institutes of Health [2R01HG007182-04A1]. The content of this paper is solely the responsibility of the authors, and does not necessarily represent the official views of our funding organizations.

## Author contributions

IB, CCH, CEC, LMNH, and TW conceived of the presented work. IB and CL designed the AMPlify model with help from CY. CL implemented the model with help from CY, FT, and RLW. IB and SAH designed the AMP discovery pipeline with help from CL. CEC, LMNH, and TW provided the bacterial strains tested. SH, CCH, CEC, LMNH, and TW developed the antimicrobial susceptibility testing protocol with input from DS, LB, and SAH. DS, LB, SAH, and SH conducted antimicrobial susceptibility testing. CL, DS, and SAH drafted the manuscript, and all authors were involved in its revision.

## Notes

### Competing Interest Statement

IB is the founder of, and a shareholder in, Amphoraxe Life Sciences Inc.

## References

1. Reardon, S. Antibiotic resistance sweeping developing world. Nature (2014). doi:10.1038/509141a

2. Brandenburg, K., Heinbockel, L., Correa, W. & Lohner, K. Peptides with dual mode of action: Killing bacteria and preventing endotoxin-induced sepsis. Biochim. Biophys. Acta - Biomembr. (2016). doi:10.1016/j.bbamem.2016.01.011

3. De Lucca, A. J. & Walsh, T. J. Antifungal peptides: Novel therapeutic compounds against emerging pathogens. Antimicrobial Agents and Chemotherapy (1999). doi:10.1128/aac.43.1.1

4. Klotman, M. E. & Chang, T. L. Defensins in innate antiviral immunity. Nature Reviews Immunology (2006). doi:10.1038/nri1860

5. Zhang, L. & Gallo, R. L. Antimicrobial peptides. Curr. Biol. (2016). doi:10.1016/j.cub.2015.11.017

6. Meylan, S., Andrews, I. W. & Collins, J. J. Targeting Antibiotic Tolerance, Pathogen by Pathogen. Cell (2018). doi:10.1016/j.cell.2018.01.037

7. Boman, H. G. Antibacterial peptides: Basic facts and emerging concepts. Journal of Internal Medicine (2003). doi:10.1046/j.1365-2796.2003.01228.x

8. Wu, Q. et al. Recent Progress in Machine Learning-based Prediction of Peptide Activity for Drug Discovery. Curr. Top. Med. Chem. (2019). doi:10.2174/1568026619666190122151634

9. Waghu, F. H., Barai, R. S., Gurung, P. & Idicula-Thomas, S. CAMPR3: A database on sequences, structures and signatures of antimicrobial peptides. Nucleic Acids Res. (2016). doi:10.1093/nar/gkv1051

10. Waghu, F. H. et al. CAMP: Collection of sequences and structures of antimicrobial peptides. Nucleic Acids Res. (2014). doi:10.1093/nar/gkt1157

11. Xiao, X., Wang, P., Lin, W. Z., Jia, J. H. & Chou, K. C. IAMP-2L: A two-level multi-label classifier for identifying antimicrobial peptides and their functional types. Anal. Biochem. (2013). doi:10.1016/j.ab.2013.01.019

12. Meher, P. K., Sahu, T. K., Saini, V. & Rao, A. R. Predicting antimicrobial peptides with improved accuracy by incorporating the compositional, physico-chemical and structural features into Chou’s general PseAAC. Sci. Rep. (2017). doi:10.1038/srep42362

13. Li, Y. et al. Deep learning in bioinformatics: Introduction, application, and perspective in the big data era. Methods (2019). doi:10.1016/j.ymeth.2019.04.008

14. Veltri, D., Kamath, U. & Shehu, A. Deep learning improves antimicrobial peptide recognition. Bioinformatics (2018). doi:10.1093/bioinformatics/bty179

15. Li, S., Li, W., Cook, C., Zhu, C. & Gao, Y. Independently Recurrent Neural Network (IndRNN): Building A Longer and Deeper RNN. In Proceedings of the IEEE Computer Society Conference on Computer Vision and Pattern Recognition (2018). doi:10.1109/CVPR.2018.00572

16. Young, T., Hazarika, D., Poria, S. & Cambria, E. Recent trends in deep learning based natural language processing [Review Article]. IEEE Computational Intelligence Magazine 13, 55–75 (2018).

17. Bahdanau, D., Cho, K. H. & Bengio, Y. Neural machine translation by jointly learning to align and translate. In 3rd International Conference on Learning Representations, ICLR 2015 - Conference Track Proceedings (2015).

18. Vaswani, A. et al. Attention is all you need. In Advances in Neural Information Processing Systems (2017).

19. Yang, Z. et al. Hierarchical attention networks for document classification. In Proceedings of the Conference of the North American Chapter of the Association for Computational Linguistics: Human Language Technologies, NAACL HLT 2016 (2016). doi:10.18653/v1/n16-1174

20. Xu, K. et al. Show, attend and tell: Neural image caption generation with visual attention. In 32nd International Conference on Machine Learning, ICML 2015 (2015).

21. Hochreiter, S. & Schmidhuber, J. Long Short-Term Memory. Neural Comput. 9, 1735–1780 (1997).

22. Gers, F. A., Schmidhuber, J. & Cummins, F. Learning to forget: Continual prediction with LSTM. Neural Comput. (2000). doi:10.1162/089976600300015015

23. Schuster, M. & Paliwal, K. K. Bidirectional recurrent neural networks. IEEE Trans. Signal Process. (1997). doi:10.1109/78.650093

24. World Health Organization. WHO publishes list of bacteria for which new antibiotics are urgently needed. (2017). Available at: https://www.who.int/en/news-room/detail/27-02-2017-who-publishes-list-of-bacteria-for-which-new-antibiotics-are-urgently-needed.

25. Bingen, E. et al. Resistance to macrolides in Streptococcus pyogenes in France in pediatric patients. Antimicrob. Agents Chemother. (2000). doi:10.1128/AAC.44.6.1453-1457.2000

26. Vanhoye, D., Bruston, F., Nicolas, P. & Amiche, M. Antimicrobial peptides from hylid and ranin frogs originated from a 150-million-year-old ancestral precursor with a conserved signal peptide but a hypermutable antimicrobial domain. Eur. J. Biochem. (2003). doi:10.1046/j.1432-1033.2003.03584.x

27. Hammond, S. A. et al. The North American bullfrog draft genome provides insight into hormonal regulation of long noncoding RNA. Nat. Commun. (2017). doi:10.1038/s41467-017-01316-7

28. Helbing, C. C. et al. Antimicrobial peptides from Rana [Lithobates] catesbeiana: Gene structure and bioinformatic identification of novel forms from tadpoles. Sci. Rep. (2019). doi:10.1038/s41598-018-38442-1

29. Zhao, R.-L., Han, J.-Y., Han, W.-Y., He, H.-X. & Ma, J.-F. Effects of Two Novel Peptides from Skin of Lithobates Catesbeianus on Tumor Cell Morphology and Proliferation. In Molecular Cloning - Selected Applications in Medicine and Biology (2011). doi:10.5772/25209

30. Novković, M., Simunić, J., Bojović, V., Tossi, A. & Juretić, D. DADP: The database of anuran defense peptides. Bioinformatics (2012). doi:10.1093/bioinformatics/bts141

31. Cameron, C. E., Brouwer, N. L., Tisch, L. M. & Kuroiwa, J. M. Y. Defining the interaction of the Treponema pallidum adhesin Tp0751 with laminin. Infect. Immun. (2005). doi:10.1128/IAI.73.11.7485-7494.2005

32. Wang, G., Li, X. & Wang, Z. APD3: The antimicrobial peptide database as a tool for research and education. Nucleic Acids Res. (2016). doi:10.1093/nar/gkv1278

33. The UniProt Consortium. UniProt: A worldwide hub of protein knowledge. Nucleic Acids Res. 47, D506–D515 (2019).

34. Bals, R. Epithelial antimicrobial peptides in host defense against infection. Respiratory Research (2000). doi:10.1186/rr25

35. Chollet, F. Keras. (2015). Available at: https://keras.io.

36. Abadi, M. et al. TensorFlow: Large-scale machine learning on heterogeneous systems. (2015). Available at: https://www.tensorflow.org.

37. Kingma, D. P. & Ba, J. L. Adam: A method for stochastic optimization. In 3rd International Conference on Learning Representations, ICLR 2015 - Conference Track Proceedings (2015).

38. Srivastava, N., Hinton, G., Krizhevsky, A., Sutskever, I. & Salakhutdinov, R. Dropout: A simple way to prevent neural networks from overfitting. J. Mach. Learn. Res. (2014).

39. Wu, T. D. & Nacu, S. Fast and SNP-tolerant detection of complex variants and splicing in short reads. Bioinformatics (2010). doi:10.1093/bioinformatics/btq057

40. Holt, C. & Yandell, M. MAKER2: An annotation pipeline and genome-database management tool for second-generation genome projects. BMC Bioinformatics (2011). doi:10.1186/1471-2105-12-491

41. Keller, O., Kollmar, M., Stanke, M. & Waack, S. A novel hybrid gene prediction method employing protein multiple sequence alignments. Bioinformatics (2011). doi:10.1093/bioinformatics/btr010

42. Slater, G. S. C. & Birney, E. Automated generation of heuristics for biological sequence comparison. BMC Bioinformatics (2005). doi:10.1186/1471-2105-6-31

43. Lomsadze, A., Ter-Hovhannisyan, V., Chernoff, Y. O. & Borodovsky, M. Gene identification in novel eukaryotic genomes by self-training algorithm. Nucleic Acids Res. (2005). doi:10.1093/nar/gki937

44. Korf, I. Gene finding in novel genomes. BMC Bioinformatics (2004). doi:10.1186/1471-2105-5-59

45. Camacho, C. et al. BLAST+: Architecture and applications. BMC Bioinformatics (2009). doi:10.1186/1471-2105-10-421

46. Smit, A., Hubley, R. & Grenn, P. RepeatMasker Open-4.0. (2015). Available at: http://www.repeatmasker.org.

47. Clinical and Laboratory Standards Institute. Methods for dilution antimicrobial susceptibility tests for bacteria that grow aerobically: approved standard. (2015).

48. Hancock, R. E. W. Modified MIC method for cationic antimicrobial peptides. (1999). Available at: http://cmdr.ubc.ca/bobh/method/modified-mic-method-for-cationic-antimicrobial-peptides/. (Accessed: 22nd September 2017)

49. Nguyen, L. T. et al. Serum stabilities of short tryptophan- and arginine-rich antimicrobial peptide analogs. PLoS One (2010). doi:10.1371/journal.pone.0012684

