## Supplementary material for "AMPlify: attentive deep learning model for discovery of novel antimicrobial peptides effective against WHO priority pathogens"

### Supplementary Notes

#### Supplementary Note 1: Performance comparison of AMP Scanner Vr.2 and AMPlify re-trained on the AMP Scanner Vr.2 dataset

In order to test whether the AMPlify architecture can still perform well on other datasets, we conducted an analysis on the AMP Scanner Vr.2 datasets[1] (named "Train", "Tune" and "Test" partitions) in two different manners. First, AMPlify was cross-validated on all data provided by AMP Scanner Vr.2 in the same way they did in their study[1]. Second, AMPlify was re-trained on the same dataset (i.e. "Train+Tune" partitions) used in the original AMP Scanner Vr.2 publication for performance comparison on the "Test" partition[1]. The results for AMP Scanner Vr.2 were taken directly from Table 1 of Veltri and co-workers' work[1], with the best tuned hyperparameters. They did not report the F1 score, so this metric is not compared here.

Supplementary Table 2 shows the 10-fold cross-validation results of AMP Scanner Vr.2 and AMPlify on all data provided by AMP Scanner Vr.2. A single sub-model of AMPlify without ensemble learning still outperforms AMP Scanner Vr.2 in accuracy (92.15% vs. 91.51%), sensitivity (91.06% vs. 88.81%) and AUROC (97.15% vs. 96.58%). Although its specificity is slightly lower, the difference is < 1%. After ensemble learning is adopted, AMPlify outperforms AMP Scanner Vr.2 in all four metrics by at least 1%.

Supplementary Table 3 shows the performance comparison on the AMP Scanner Vr.2 test set between AMP scanner Vr.2 and the re-trained version (on "Train+Tune" partitions) of AMPlify. AMPlify re-trained on their dataset still outperforms AMP Scanner Vr.2 in all four metrics of accuracy (92.35% vs. 91.01%), sensitivity (90.59% vs. 89.89%), specificity (94.10% vs. 92.13%) and AUROC (97.00% vs. 96.48%). However, each sub-model of AMPlify re-trained on their dataset does not show substantial advantage in overall accuracy over AMP Scanner Vr.2. The first reason is that AMPlify is trained on fewer data than AMP Scanner Vr.2 due to the early stopping technique used. The second reason is that the validation sets we set aside for early stopping here are too small compared with those used in our own training process (because the entire set for training here is only 1/3 of ours in size). Note that the 10-fold cross-validation does not suffer from such problems because it is done on all data, with both the training and the early stopping monitored validation sets of single sub-model within each fold larger. After a train-validation split for early stopping, the new training and early stopping monitored validation sets for test comparison are comprised of only 1704–1706 and 426–428 sequences respectively, while those for 10-fold cross-validation are 2560–2562 and 640–642 sequences respectively. Supplementary Fig. 1 shows a comparison between the learning curves of five sub-models trained on our own training set and on their original dataset, respectively. Obviously, the validation accuracy fluctuates greatly when the models are trained on their dataset, which suggests that those validation sets are too small to represent the distribution of the entire dataset. Hence, sub-models of AMPlify are not well trained on their dataset since it is difficult for early stopping to decide the best epochs in such cases. Both of the above reasons conclude that there is a lack of sufficient data in their training sets to train and validate AMPlify.

In summary, AMPlify and its sub-models can outperform the state-of-the-art AMP Scanner Vr.2 with respect to the 10-fold cross-validation analysis on their dataset. Further, AMPlify re-trained on their dataset is still able to outperform AMP Scanner Vr.2, and those sub-models of AMPlify are likely to gain better performance if the training set is large enough.

### Supplementary Figures

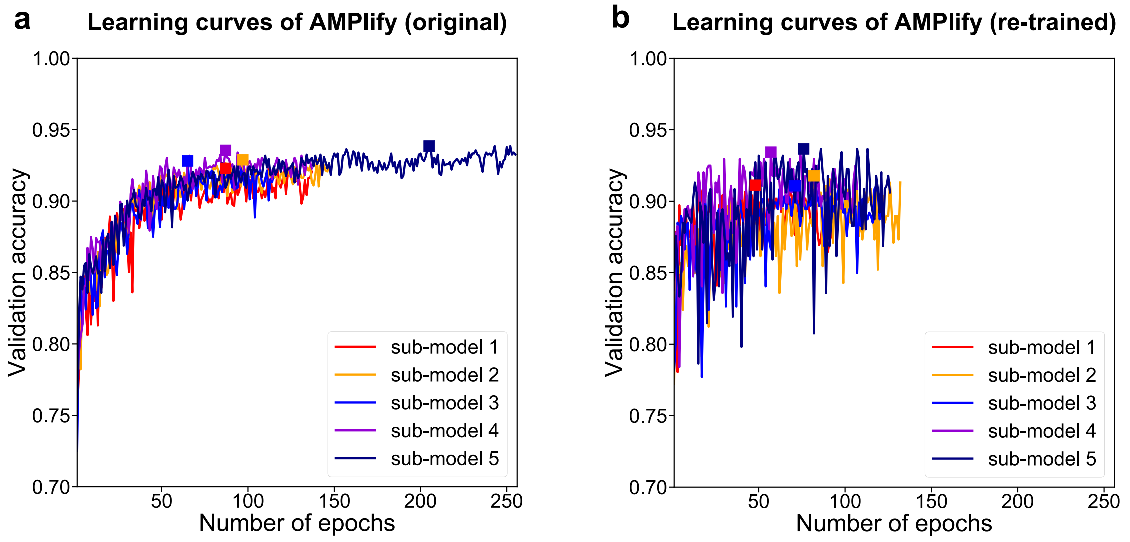


**Supplementary Figure 1: Learning curve comparison (on the validation sets for early stopping) of sub-models of AMPlify trained on two different datasets. (a)** Sub-models of AMPlify trained on our own training set; **(b)** Sub-models of AMPlify trained on the AMP Scanner Vr.2 "Train+Tune" partitions. Square markers denote the best epochs chosen by early stopping, and the x-axes have been set in the same range in order for a clearer comparison.

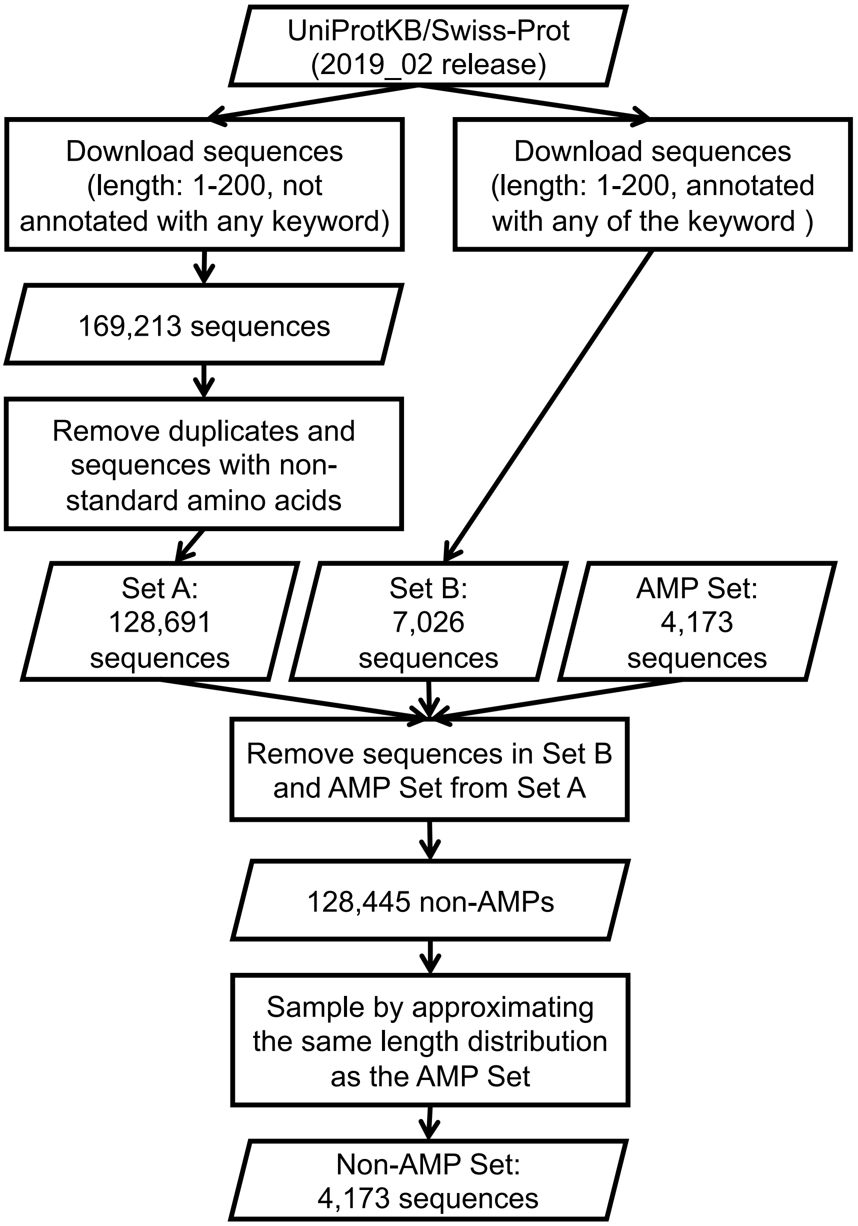


**Supplementary Figure 2: Workflow of the non-AMP sampling process.** This workflow shows how we obtained 4,173 non-AMP sequences from the UniProtKB/Swiss-Prot database.

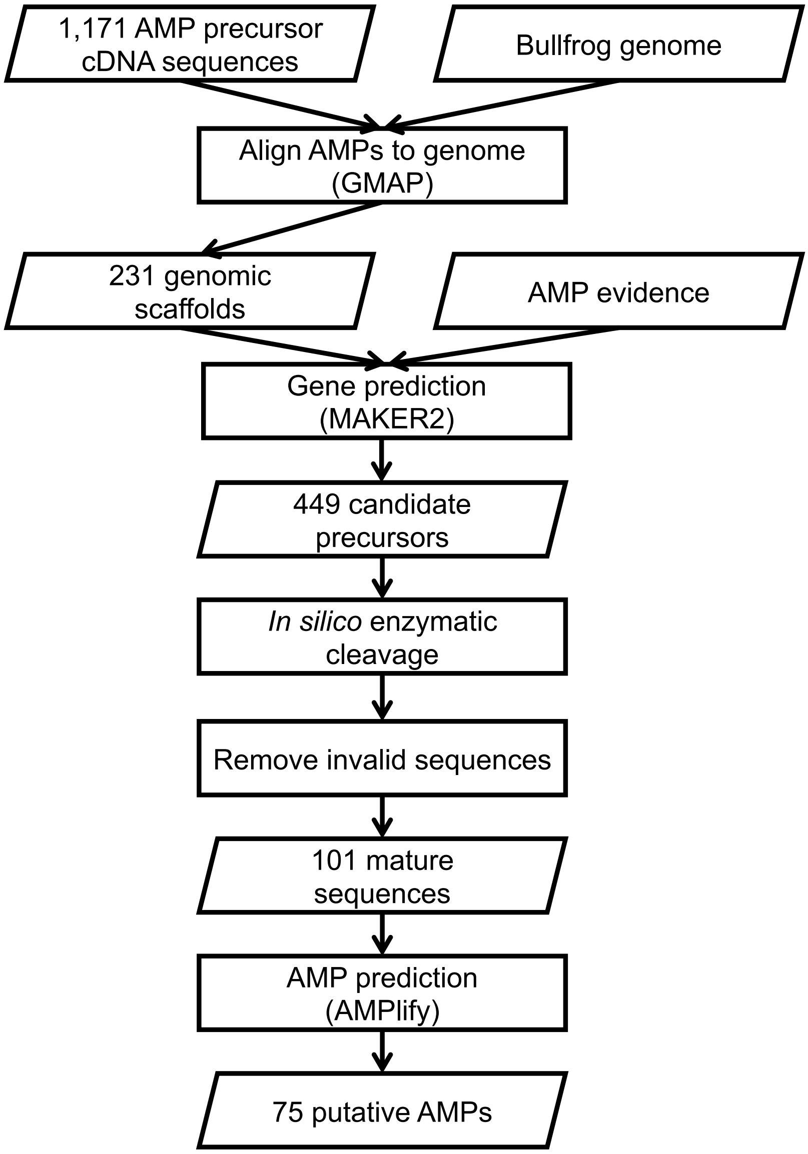


**Supplementary Figure 3: An overview of the AMP discovery pipeline we employed.** This workflow shows how we identified putative AMPs from the bullfrog genome. Invalid sequences denote those not suitable for AMPlify prediction (with lengths not between 2 and 200 or with non-standard amino acids).

### Supplementary Tables

**Supplementary Table 1: Stratified 5-fold cross-validation results of different architectures on the training set.** The top section compares the architecture of AMPlify, with and without ensemble learning, with its simpler variations. The second section shows the architecture of AMP Scanner Vr.2 cross-validated on our training set. Values of accuracy (acc), sensitivity (sens), specificity (spec), F1 score (F1) and area under the receiver operating characteristic curve (AUROC) are presented along with their standard deviations in percentage.

| Model Architecture | Acc | Sens | Spec | F1 | AUROC |
|---|---|---|---|---|---|
| Bi-LSTM | 89.04±1.02 | 89.10±1.14 | 88.98±1.80 | 89.05±0.98 | 95.82±0.56 |
| Bi-LSTM + CA | 90.13±1.02 | 90.05±0.94 | 90.20±1.91 | 90.13±0.95 | 96.14±0.37 |
| Bi-LSTM + MHSDPA + CA | 91.70±0.66 | 91.40±0.71 | 92.00±1.51 | 91.68±0.61 | 96.92±0.30 |
| **AMPlify**: Bi-LSTM + MHSDPA + CA (with ensemble) | **92.79±0.77** | **92.12±1.28** | **93.47±0.53** | **92.74±0.80** | **97.44±0.53** |
| **AMP Scanner Vr.2:** Embedding + Conv + Max Pooling + LSTM | | | | | |
| - [10 epochs[a]] | 90.55±1.43 | 90.14±1.62 | 90.95±4.08 | 90.54±1.17 | 96.40±0.58 |
| - [early stopped[b]] | 90.52±0.76 | 89.45±1.10 | 91.58±1.48 | 90.42±0.74 | 96.35±0.43 |

[a]The best hyperparameter as stated in the paper.
[b]Optimal numbers of training epochs determined by early stopping range from 17 to 71.

**Supplementary Table 2: Comparison between AMP Scanner Vr.2 and AMPlify cross-validated on all data provided by AMP Scanner Vr.2 ("Train+Tune+Test" partitions).** This table shows the 10-fold cross-validation results of AMP Scanner Vr.2 and AMPlify on all data provided by AMP Scanner Vr.2. Values of accuracy (acc), sensitivity (sens), specificity (spec) and area under the receiver operating characteristic curve (AUROC) are presented along with their standard deviations in percentage.

| Model | Acc | Sens | Spec | AUROC |
|---|---|---|---|---|
| AMP Scanner Vr.2 | 91.51±0.89 | 88.81±3.53 | 94.21±2.68 | 96.58±0.66 |
| AMPlify (re-trained, single model) | 92.15±0.86 | **91.06±1.76** | 93.25±2.51 | 97.15±0.58 |
| AMPlify (re-trained, ensemble) | **93.22±0.71** | 90.55±1.62 | **95.89±1.56** | **97.61±0.46** |

**Supplementary Table 3: Performance comparison between AMP Scanner Vr.2 and AMPlify re-trained on the AMP Scanner Vr.2 "Train+Tune" partitions and tested on their "Test" partition.** Since AMPlify applies early stopping and the exact size of training set for each sub-model is smaller, the exact training size for each model is listed here in the second column. Values of accuracy (acc), sensitivity (sens), specificity (spec) and area under the receiver operating characteristic curve (AUROC) are presented in percentage.

| **Model** | **Training set size** | **Acc** | **Sens** | **Spec** | **AUROC** |
|---|---|---|---|---|---|
| AMP Scanner Vr.2 | 2132 | 91.01 | 89.89 | 92.13 | 96.48 |
| AMPlify (re-trained) | 2132 | **92.35** | 90.59 | 94.10 | **97.00** |
| - [sub-model 1] | 1704 | 91.01 | **91.85** | 90.17 | 96.29 |
| - [sub-model 2] | 1706 | 91.36 | 88.76 | 93.96 | 96.21 |
| - [sub-model 3] | 1706 | 91.71 | 91.43 | 91.99 | 96.74 |
| - [sub-model 4] | 1706 | 91.22 | 88.06 | 94.38 | 96.14 |
| - [sub-model 5] | 1706 | 91.36 | 88.20 | **94.52** | 96.20 |

#### List of Abbreviations

acc: Accuracy

AUROC: Area under the receiver operating characteristic curve

Bi-LSTM: Bidirectional long short-term memory

CA: Context attention

Conv: Convolutional

LSTM: Long short-term memory

MHSDPA: Multi-head scaled dot-product attention

sens: Sensitivity

spec: Specificity

#### Reference


1. Veltri, D., Kamath, U. & Shehu, A. Deep learning improves antimicrobial peptide recognition. *Bioinformatics* (2018). doi:10.1093/bioinformatics/bty179
